# Hyperflexion is unlikely to be the primary cervical spine injury mechanism in accidental head-on rugby tackling

**DOI:** 10.1101/2022.02.18.481008

**Authors:** Pavlos Silvestros, Ezio Preatoni, Harinderjit S. Gill, Dario Cazzola

**Author notes:** <u>Corresponding Author</u>, Dario Cazzola, 1.309 Applied Biomechanics Suite, University of Bath, Bath, BA2 7AY, United Kingdom, <u>e-mail</u>, <u>telephone number</u>.

## Abstract

In Rugby a high proportion of catastrophic cervical spine injuries occur during tackling. In the injury prevention literature, there is still an open debate on the injury mechanisms related to such injuries, with hyperflexion and buckling being under scrutiny. The aims of this study were to determine the primary cervical spine injury mechanism during head-on rugby tackling, and evaluate the effect of tackling technique on cervical spine intervertebral loading. We conducted an *in silico* study to examine the dynamic response of the cervical spine under loading conditions representative of accidental head-on rugby tackles by using a subject-specific musculoskeletal model of a rugby player. The computer simulations were driven by experimental *in vivo* data of an academy rugby player tackling a punchbag, and *in vitro* data of head-first impacts using a dummy head. Results showed that: i) the earlier generation of high compression and anterior shear loads with low values of flexion moments provides evidence that hyperflexion is unlikely to be the primary injury mechanism in the sub-axial cervical spine (C3-C7) during central and posterior head impact locations; ii) a higher degree of neck flexion at impact poses the cervical spine in a more hazardous position. These findings provide objective evidence to inform injury prevention strategies or rugby law changes, with the final view of improving the safety of the game of rugby.

## Introduction

Rugby Union is a full contact field sport with the tackle, a major component of the game, resulting in a high proportion of head, neck and shoulder injuries ^1–3^. Over the last decade, epidemiological and biomechanical injury prevention research has primarily focused on mild traumatic head injuries (mTBI), and suggested that changes in tackling technique ^3–5^ could reduce the risk of concussion. However, tackling in rugby is not only associated with mTBI, but it also carries with it a high proportion (>30%) of all catastrophic cervical spine injuries in rugby^6^. Although the likelihood of sustaining a neck injury is rare (4.5/1000 playing hours)^7,8^ compared to that of concussion in the tackle (8.9/1000 playing hours)^9^, the former has higher reduction on quality of life and associated financial costs^10^. Consequently, it is fundamental to avoid the situation in which a specific policy change is beneficial for one type of injury (e.g. mTBI), and detrimental for other more severe ones^11^ (e.g. cervical spine injuries). Thus, there is a pressing need to investigate neck injury mechanisms and their relationship with technique changes, to appropriately inform injury prevention interventions to increase the safety of the game of rugby^12^.

The main theorised cervical spine injury mechanisms in rugby injury prevention research are buckling ^13^ and hyperflexion ^14^. Buckling is caused by a compressive axial load applied to the cervical spine column that results in the combination of flexion and extension across the intervertebral joints with complex loading patterns that includes bending moments, compressive and shear forces^15,16^. Hyperflexion is the rapid posterior to anterior head motion resulting in intervertebral joints exceeding their physiological flexion range. The catastrophic cervical spine dislocations observed in rugby accidents are predominately anterior bilateral facet dislocations in the lower cervical spine (C4-C5 to C6-C7) ^13^. Hyperflexion was deemed the primary injury mechanism for anterior cervical spine dislocations in rugby by Dennison et al. (2012) based on uncertainty regarding the buckling mechanism^14^, but this theory is only supported by player recollections ^17^ and video analysis of the inciting events. Additionally, despite the lack of more objective biomechanical analyses, such evidence was used to draw a cause-effect relationship with clinically observed spinal injuries. On the other hand, cervical spine buckling had been recreated during quasi-static and dynamic *in vitro* cadaveric experiments ^15,18–20^.

These experimental studies showed that cervical spine buckling is sensitive to neck pre-flexion angles (geometric alignment), simulated muscle forces (internal stability), impact load characteristics and interaction with impacted surface (endpoint constraint) ^15,18,21–23^. The rationale for questioning buckling as the primary injury mechanism in rugby cervical spine injuries was, firstly, that highly controlled *in vitro* experiments differ greatly from the real *in-vivo* dynamics of rugby tackles. Secondly, qualitative data from video analysis and personal accounts of injured players supported hyperflexion as the mechanism of injury. Computer simulations have since proven a valuable method in being able to recreate with high fidelity the internal (i.e. muscle and joint contact forces) and external loading conditions during which cervical spine injuries occur under inertial and compressive loading ^24–28^. *In-silico* simulations using musculoskeletal models have strengthened the theory that muscle forces affect resulting head and neck dynamics during injurious scenarios (inertial and axial impacts) ^24,28,29^. However, arbitrary levels of simulated muscle activations and resulting muscle forces have been applied in these studies, which limit their representability of the event under investigation.

Biomechanical models of the neck validated for dynamic loading have been able to characterise the internal loading patterns and resulting kinematics of cervical vertebrae during impacts which is not achievable *in vitro* and *in vivo.* Computational investigations ^28^ have supported the theorised decoupling between externally observed head and neck kinematics and the internal dynamic response of the spine during axial loading injuries ^15,16,19^. These have supported buckling over hyperflexion as the main injury mechanism under compressive impacts to the head ^23,28^. However, these studies were not focused on rugby tackling, and such evidences has not been translated into injury prevention practices in relation to minimising cervical spine injuries occurrence.

A rugby tackle is a very dynamic sporting event, characterised by extremely variable and intense external loading conditions and a high level of spinal muscle co-contraction ^30^. It is also very challenging to measure accurate rugby tackling forces *in vivo* to inform and drive computational studies. For these reasons, a rugby-specific computational study has not yet been conducted to unveil the predominant cervical spine injury mechanism observed during tackling. Therefore, a rugby-specific theoretical study should aim to replicate high risk impact scenarios associated with catastrophic neck injuries occurring in gameplay situations and relate them to applied aspects including player tackling technique and governing laws of the game.

To address this gap in the literature and inform injury prevention practices we conducted an *in silico* investigation, where a combination of *in vitro* and *in vivo* data were used to examine the dynamic response of the cervical spine to loading conditions representative of head-on rugby tackles. The aims of the study were firstly to determine the primary cervical spine injury mechanism during accidental head-on rugby tackling, and secondly to highlight the effect of tackling technique on the intervertebral loading experienced during high fidelity musculoskeletal simulations.

## Methods

### Experimental data

#### In vivo data collection

One academy-level front-row rugby player (male, 22 years, 1.82 m, 113.7 kg) participated in this study. Ethical approval was obtained from the Research Ethics Approval Committee for Health of the University of Bath (approval number: EP 15/16 131) and the participant provided written informed consent prior to data collection. The experimental data collection on human participant was performed in accordance with the Declaration of Helsinki. Full body kinematics (Oqus, Qualysis, Sweden) and bilateral electromyography (EMG) recordings (Trigno, Delsys, USA) of the sternocleidomastoid and upper trapezius muscles were collected at 250 Hz and 2500 Hz, respectively during laboratory-based staged tackling trials with a tackle simulator ^30,31^. The EMG signals were band-pass filtered (10-250 Hz; maintaining 97% of signal power), full wave rectified, low-pass filtered at 6 Hz^32^ with the same filter, then amplitude normalised to the maximum recorded value identified in maximum voluntary contraction trials (Table 1) to create EMG linear envelopes in Matlab R2017a (The Mathworks Inc., Natick MA, USA). Kinematics and EMG signals at the instant of tackle impact were used to inform the initial conditions of the model during the *in silico* simulations. High resolution isotropic (1 mm^3^ voxel) T1-weighted MRI (Skyra, SEIMENS, Germany) imaging sequences were collected from the participant on the same day as experimental data collection. The MRI sequences were semi-automatically segmented and used to create a personalised musculoskeletal model^33^ as described in the following section of this study.

**Table 1.**
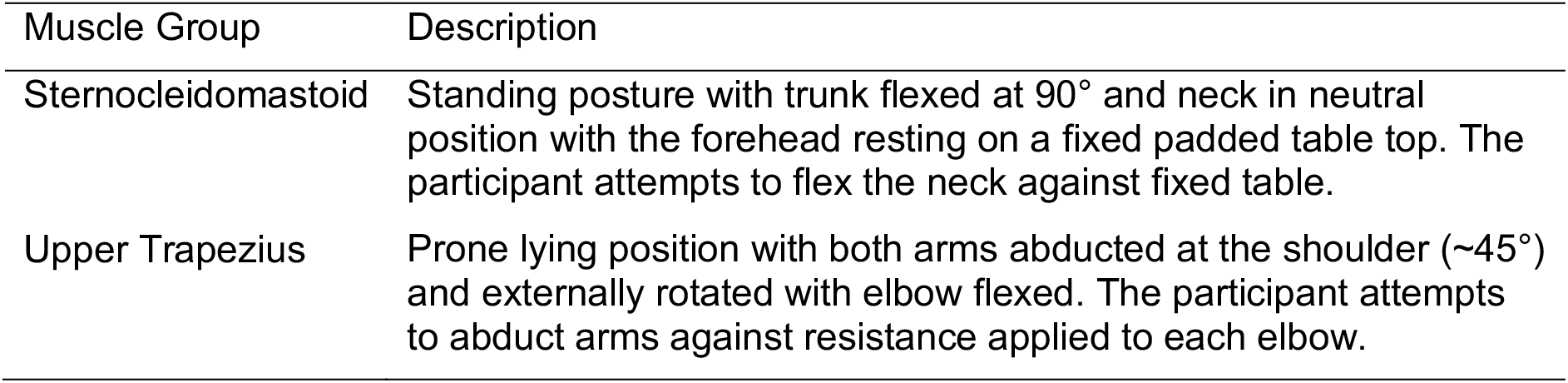
Positions used to establish maximum voluntary contraction for the sternocleidomastoid and upper trapezius muscle groups

#### In vitro data collection

A head and neck assembly of an anthropometric test device (ATD) (Hybrid III 50^th^ percentile male, Human Kinetics, Germany) was attached to a steel frame 1.5 m from a ground anchoring and used to simulate accidental head-on rugby tackle impacts to the head. A six-axis load cell was instrumented at the head and neck interface of the ATD to measure forces caused by the impacts of the tackle simulator with the ATD assembly. Impacts were generated by the tackle simulator contacting the ATD assembly at two different speeds 2.0-2.5 m/s and 3.1-3.6 m/s^30,34^. These impacts aimed to represent the momentum change experienced during live tackles ^35^. The resultant impact force magnitudes and loading rates were used to inform theoretical impact conditions applied during the *in silico* simulations.

### Musculoskeletal simulations

#### Musculoskeletal Model

The population specific male-forward Rugby Model^31^ was updated and used as the baseline model for this study. The main updates were carried out in OpenSim 3.3^36^ and consisted of the: i) inclusion of hyoid muscle group to improve its physiological fidelity^37^, ii) personalisation of cervical vertebrae dimensions using MRI images; iii) integration of wrapping surfaces to better replicate Muscle-tendon units (MTU) lines of action in the cervical spine.

More specifically, the neck region (C1-C7) of the musculoskeletal model was scaled in each dimension (height, width and depth) based on anatomical measurements of the participant’s cervical vertebrae from the segmented MRI images. MTU attachment sites were not changed with respect to the population specific model^31^ (taken from Vasavada et al.,1998)^38^, due to difficulties in identifying muscle attachment locations in the MRI. The remaining model segments were linearly scaled based on anatomical motion capture markers. The dynamics of each sub-axial cervical spine joint (C2-C3 to C6-C7) was modelled using validated six degree-of-freedom (6 DOF) Kelvin-Voigt bushings^39^. These bushings characterised the dynamic response of intervertebral cervical joints under high energy impacts.

The wrapping surfaces were defined using the MRI scans of the participant tested and were added to the musculoskeletal model. The wrapping surfaces included: i) a cylinder anterior to the lower cervical spine registered to the C6 vertebra^40^; ii) a sphere originating and registered to the C2 vertebra; iii) two bilateral cylinders at the posterior of the upper cervical spine also registered to the C2 vertebra; iv) lastly two bilateral tori at the lower cervical spine registered to the C7 vertebra. All wrapping surfaces were constrained to move with their registered bodies as explained in Silvestros et al.^33^. The newly developed model is available from the SimTK repository (https://simtk.org/projects/csibath).

#### Neck angle conditions

To examine the effect of initial neck positioning on intervertebral loading during impacts in head-on tackles the model’s sub-axial neck angles (C2-C3 to C7-T1) about the three axes of rotation (Flexion/Extension; Lateral Bending and Axial Rotation) were compared in steps of 5°. Specifically, the step limits for each axis were: in the sagittal plane: from 30° degrees of extension to −30° of flexion (13 conditions), frontal plane: from 0° (neutral) to −10 of lateral bending (3 conditions) and in the transverse plane: 5° to 15° of axial rotation (3 conditions). This resulted in 117 unique initial neck angle configurations. The angle ranges selected were informed by kinematic measurements of one-on-one experimental tackling trials of university/professional level rugby players ^41^. The initial angle of the upper cervical spine (C0-C1 to C1-C2) of the model was initialised the same as the *in vivo* experimental trials, which was in the extended position (18°) to replicate a more head-up position of the tackler which is widely coached for improved tackle technique (rugbysmart.co.nz)^42,43^. Neck and head angular velocities from an *in vivo* experimental tackling trial at the moment of tackle impact were also prescribed to the model for all unique initial neck angle configurations. After the initialisation of the model (t=0) the head and neck kinematics were not constrained in any position and followed the dynamics of the model.

#### Muscle activations

For all the simulations the model’s muscles were prescribed the same activation scheme (Figure 2). This activation scheme was estimated using an EMG-assisted neuromusculoskeletal model^33^ to minimise the error between experimental and simulated joint moments and muscle activations during the same *in vivo* experimental trial used for informing initial angular velocities. The estimation of the model’s 96 muscle activation patterns was solved using the Calibrated EMG-Informed Neuromusculoskeletal Modelling (CEINMS) OpenSim Toolbox^33,44,45^ that minimised the following cost function (Equation 1):

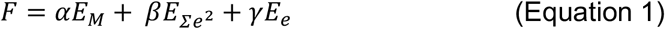

Where *E_M_* was the sum of the squared differences between the estimated and net joint moments from the inverse dynamics (sagittal and frontal plane moments of the C0-C1 through to C6-C7 joints), *E_∑e^2^_* was the sum of the squared synthesised activations for all MTUs, and *E_e_* was the sum of the differences between the adjusted model activations and experimental activations. Factors *α*, *β* and *γ* were non-negative weightings for each term of the cost function. Activation dynamics were characterised by a critically damped linear second-order differential system^32,44^. It was assumed that the MTU tendons of the model were stiff due to their short length and function in the neck.

This provides a reasonable and physiologically plausible muscle recruitment pattern and subsequent internal muscle forces applied to the cervical spine for a tackler expecting to receive contact to the shoulder when completing a wrap tackle. This aimed to approximate the conditions when a tackler is preparing to complete a wrap tackle with correct technique but however receives an accidental “head-on” impact. An accidental “head-on” impact is thus defined as the incorrect tackle technique that is not a normal phase of play where the tackler misaligns the position of their head, neck and torso resulting in the contact of their head with the oncoming attacking player. Muscle activation values were selected from instant of tackle impact during the staged tackle trial then applied to the musculoskeletal model as preactivation and remained constant for the duration of the 50 ms forward dynamic simulations. Constant activations were selected to represent muscle pre-activation as cervical spine reflex times exceed 50-60 ms ^46–48^ reducing the effect of active neck muscle modulation during short impact events.

#### Loading conditions

To replicate the possible head impact locations during accidental head-on tackles seven loading conditions were defined for the simulations (Figure 1). As the location and direction of contact forces cannot be generated with great validity in multibody musculoskeletal models, an approximation was adopted to calculate these parameters based on the skull’s geometry in Matlab R2017a (MathWorks Inc., Natick, MA, USA). Three points of impact force application were defined on the cranial midline of the model’s skull segment. These were at the vertex, posterior to the vertex (near the skull lambda or crown) and anterior to the vertex. The directional vector of these three loading conditions was defined from the points of application to the base of the skull to simulate accidental tackles resulting in “head-on” impacts (Figure 1; CA, CC, CP). Four remaining points of impact were defined on the right lateral side of the skull with an inferolatateral direction representing more oblique impacts (Figure 1: LP; LMP; LMA; LA). All points of application and directional vectors were constant with respect to the model’s skull reference system. The magnitude and loading rate of each condition was acquired from the *in vitro* experimental trials simulating head-on rugby tackle impacts to the head using the ATD and tackle simulator at two different speeds. The loading rate of these tests (80 kN/s) were one order of magnitude lower than what the bushing elements were validated against (800 kN/s). However, it has been shown that the stiffness response of intervertebral discs does not change considerably after a rate of 75-90 N/s ^49,50^ therefore the bushings used in the model were deemed valid for the loading conditions tested.

**Figure 1.**
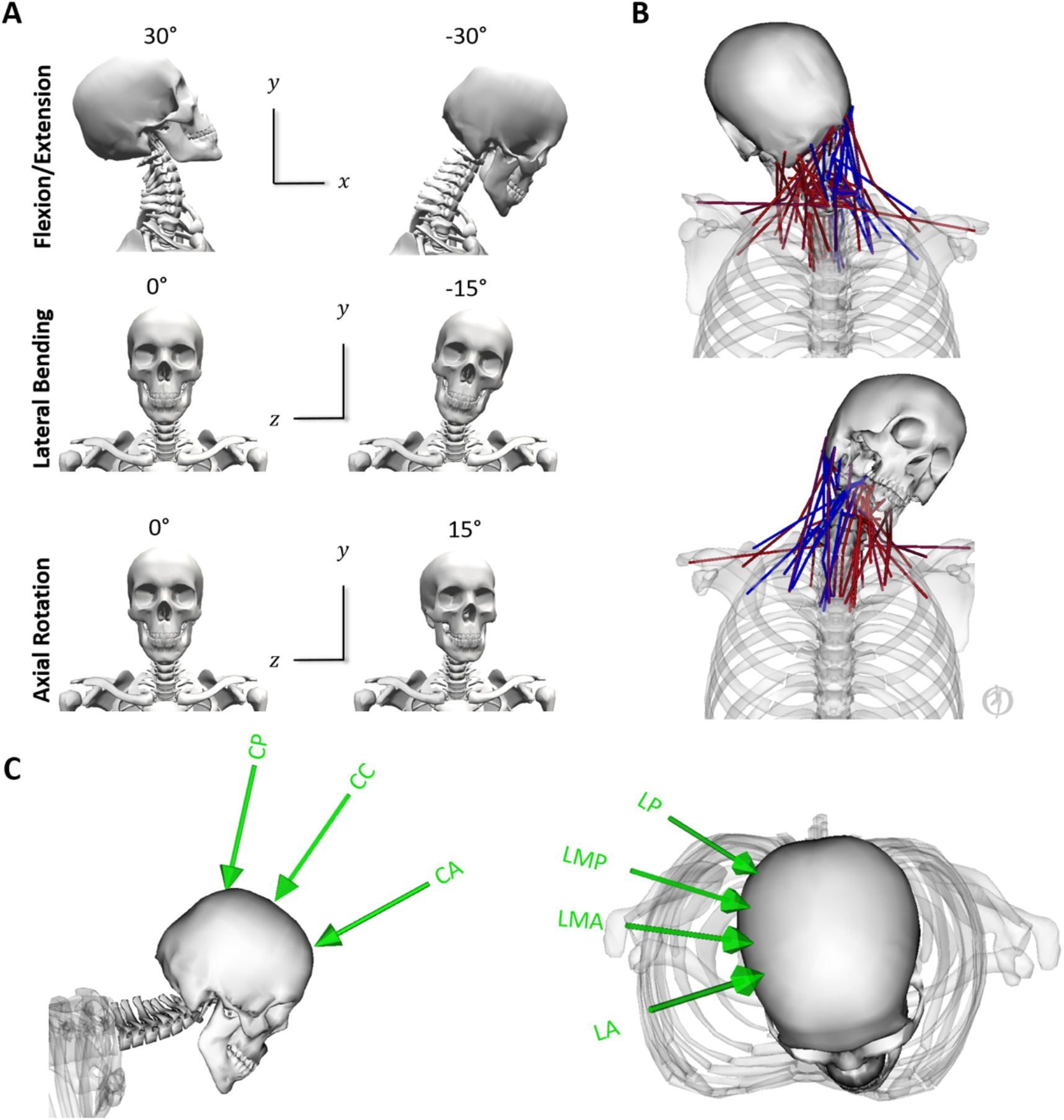
A) Close up view of the scaled MRI informed OpenSim model’s head and neck region with reference views of maximal ranges of motion tested in the simulations (muscles and wrapping surfaces removed for clarity of the cervical spine structure). Anteroposterior shear and Lateral Bending are defined by the X axis. Compression and Axial Rotation are defined by the Y axis. Lateral Shear and Flexion/Extension are defined by the Z axis. B) Neck muscle activation pattern estimated using EMG-assisted optimisation from staged experimental tackling and used across all simulations. During the staged experimental trials, the tackle was taken on the right shoulder which can be seen by the different levels of the model’s muscle activations (red – maximum, blue – minimum). C) Cranial (left) and lateral (right) loading conditions applied to the skull (CA – Cranial Anterior; CC – Cranial Central; CP – Cranial Posterior; LP – Lateral Posterior; LMP – Lateral Mid Posterior; LMA – Lateral Mid Anterior; LA – Lateral Anterior). These were identified in Matlab by defining a grid of parallel transverse (n=2) and frontal (n=5) planes at 30 mm intervals to the skull segment’s geometry. The intersecting locations of the planes with the skull’s geometry defined four rectangular regions on the right side of the skull. Each parallelogram can be thought of an impact area on the musculoskeletal model skull to which a new plane was fitted. The point of impact force application for each of the four areas defined by the rectangular regions was the projected midpoint of the fitted plane onto the skull geometry. The directional vector was the normal vector of the fitted plane directed into the skull.

#### Forward dynamics simulations

For each initial neck angle configuration, the model was loaded under the seven different loading conditions at two loading rates resulting in a total of 1,638 simulations (117 neck angles configurations x 7 loading conditions x 2 loading rates). Each simulation was performed for 50 ms from the time of initial force application and initialisation of muscle forces which were applied throughout the simulation. The 50 ms time duration for the simulations was chosen as it contained the initial measured force peak and previous *in-vitro* cadaveric head drop experiments^15^ have detected injury within this time frame. The simulations were not performed past the peak of the applied load as multibody models are unable to simulate tissue deformation and thus are not expected to reliably predict the injury in such conditions. The effects of initial neck angle and loading conditions were evaluated by analysing the maximal compressive loading, anteroposterior shear loading and flexion bending moment at the C3-C4 to C6-C7 joints sustained during the 50 ms impact simulations (Figure 2).

**Figure 2.**
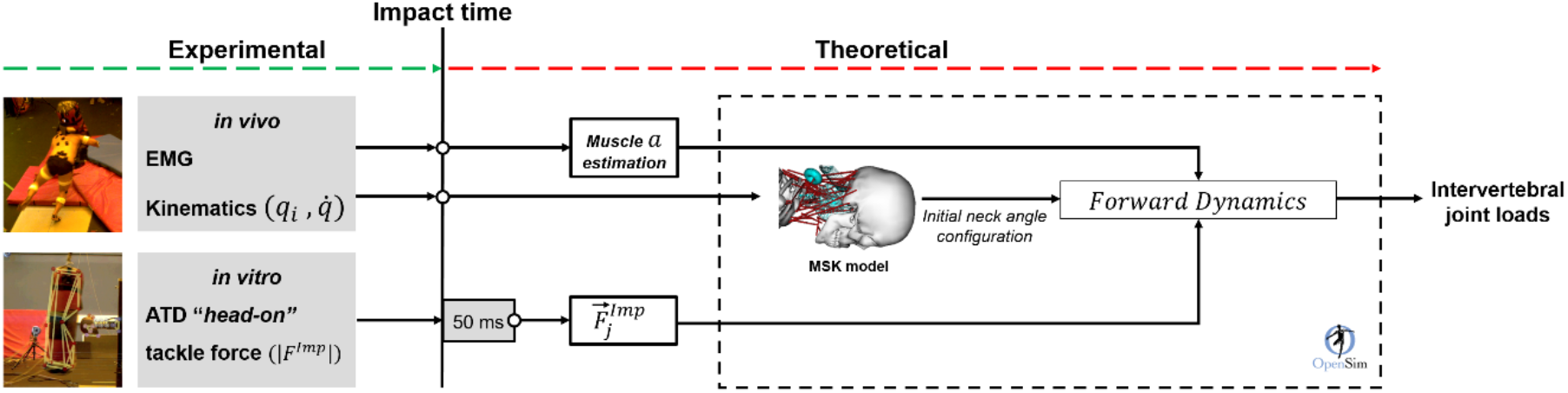
Workflow of integrated experimental and theoretical framework used to investigate cervical spine injury mechanism in rugby tackles. <u>Experimental:</u>*in vivo* data (neck muscle EMG and joint angles and velocities) were collected during stage tackling laboratory trials using a tackle simulator (mass=40 kg, velocity=3 m/s). *In vitro* data (force magnitude and loading rate) was collected from the Anthropometric Test Device (ATD) during simulated head-on impacts to the head. <u>Theoretical:</u> for each of the 1638 simulations an initial neck angle configuration combining Flexion/Extension, Lateral Bending and Axial Rotation angles (*q_i_*, n=117) taken from measured ranges in the literature^41,51^ was prescribed to the model. *In vivo* data at the time of impact were used to inform then initial neck joint angular velocities 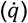 and joint angles of the torso and upper limbs. Level of neck muscle activations *(a)*at the time of impact derived from EMG-assisted analysis of the staged tackling trial were applied to the model’s muscles to be constant throughout the 50 ms simulations. For each initial neck angle configuration (*q_i_*) external loading conditions were applied (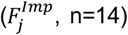, n=14) replicating different impact locations on the head at two different speeds. The points of application and direction of the loading conditions were defined using the model’s skull geometry in Matlab. The magnitude and loading rate characteristics were taken from the first 50 ms of the *in vitro* ATD impact forces.

## Results

Initial neck angles and loading conditions had the largest effect on intervertebral joint loading patterns across the cervical spine (Figure 3). Joint loads were more sensitive to initial neck flexion angles compared to changes in lateral bending and axial rotation across the loading conditions and vertebral levels (Figure 4 and 5). Average compressive joint loads were larger in the lower cervical spine whereas anteroposterior shear and flexion moments showed more complex loading patterns across intervertebral joints (Figure 6 and 7).

**Figure 3.**
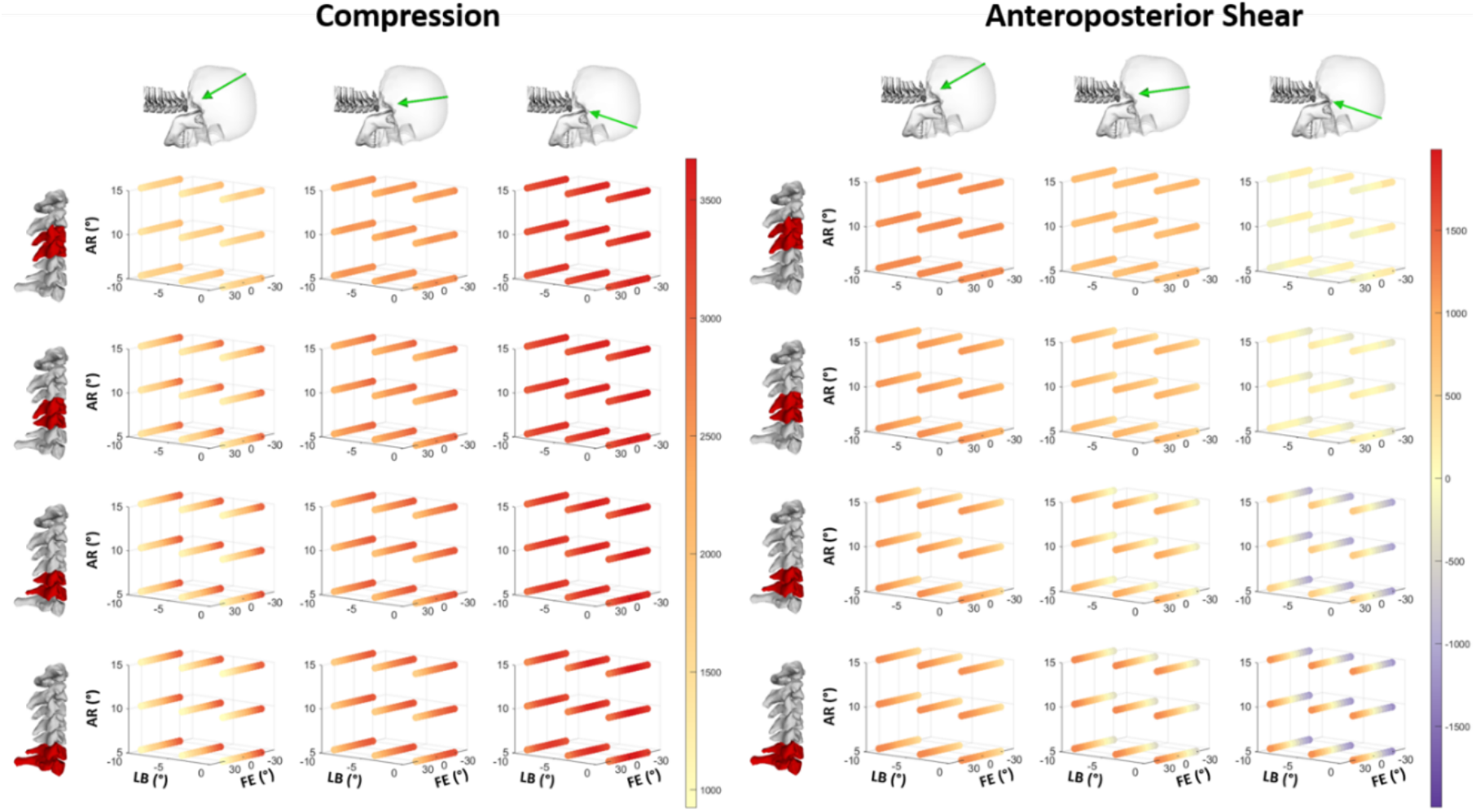
Overview presentation of the effects of neck angle (FE = flexion/extension, LB = lateral bending, and AR = axial rotation), external cranial loading location (posterior, central and anterior) on maximal cervical spine loading across the cervical spine. Patterns of maximal compression (left) and anteroposterior shear (right) loading in Newton sustained during the 50 ms simulations across all simulated initial neck angles for cranial loading conditions. Column represents an individual loading condition (Cranial Posterior – left columns; Cranial Central – centre columns and Cranial Anterior – right columns). Rows represent the cervical spine levels from C3-C4 (top) to C6-C7 (bottom). The cubic grids of each subplot represent the initial neck angle (°) in Flexion/Extension (FE), Lateral Bending (LB) and Axial Rotation (AR). Magnitude of maximal loading (newton) in the 50 ms simulations is represented with the colour bars. Note compression are only positive values and anteroposterior shear positive and negative values to represent direction with anterior and posterior respectively.

**Figure 4.**
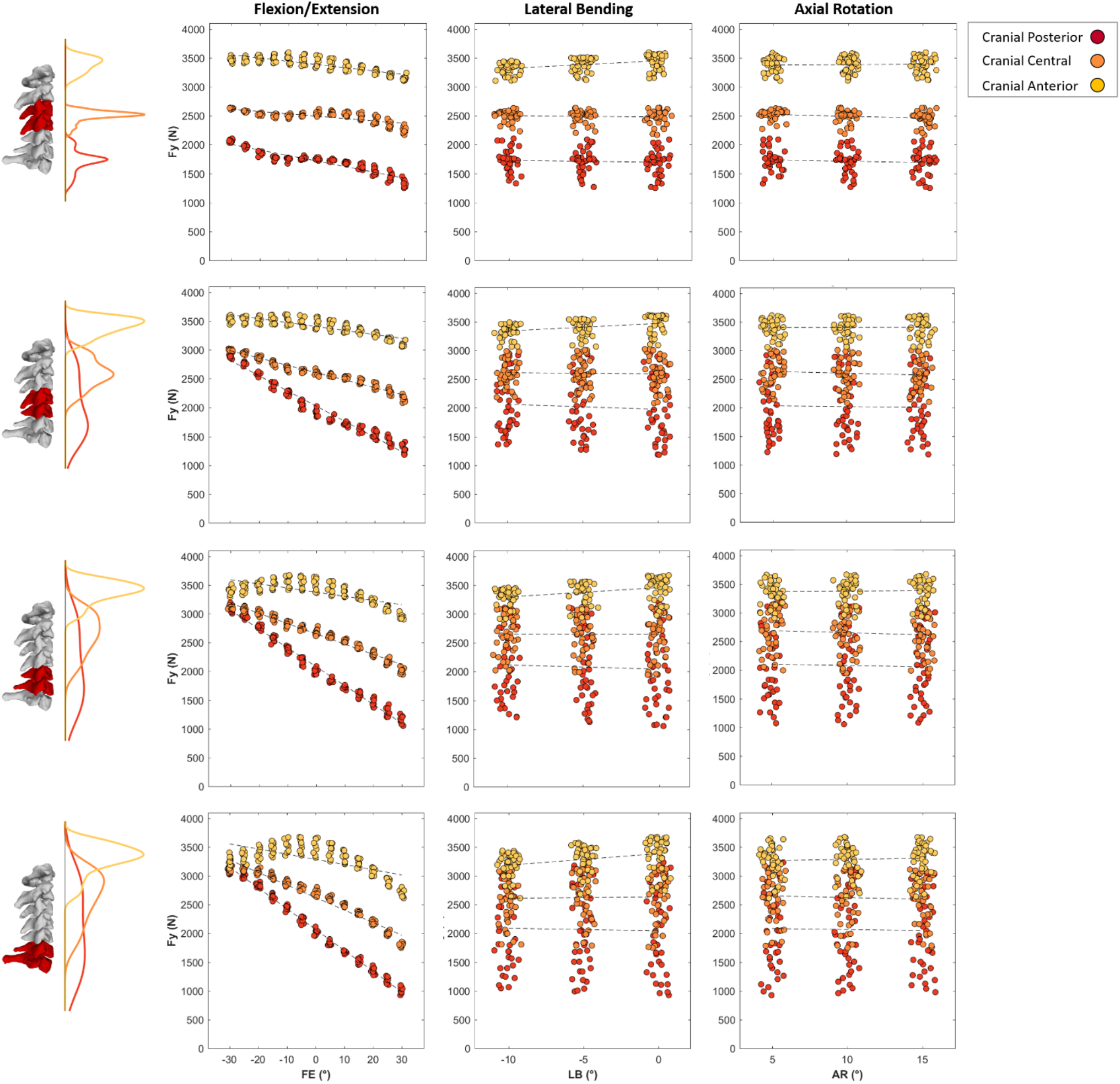
Maximal compressive joint loads (newton) of C3-C4 (top row) to C6-C7 (bottom row) intervertebral joints plotted against 5° changes in Flexion(−)/Extension(+) (left column), Lateral Bending (centre column) and Axial Rotation (right column) during the cranial loading conditions (Cranial Posterior, Cranial Central and Cranial Anterior). Kernel density estimate plots to the left of the subplot rows represent the frequency distribution density of the maximal joint loads on the vertical axes for each loading condition. First order polynomial lines of best fit are plotted to highlight the effect of joint angle on compressive joint loads for each loading condition (dashed lines). In each subplot data points are spread slightly in each 5° bin on the horizontal axes for better visualisation.

**Figure 5.**
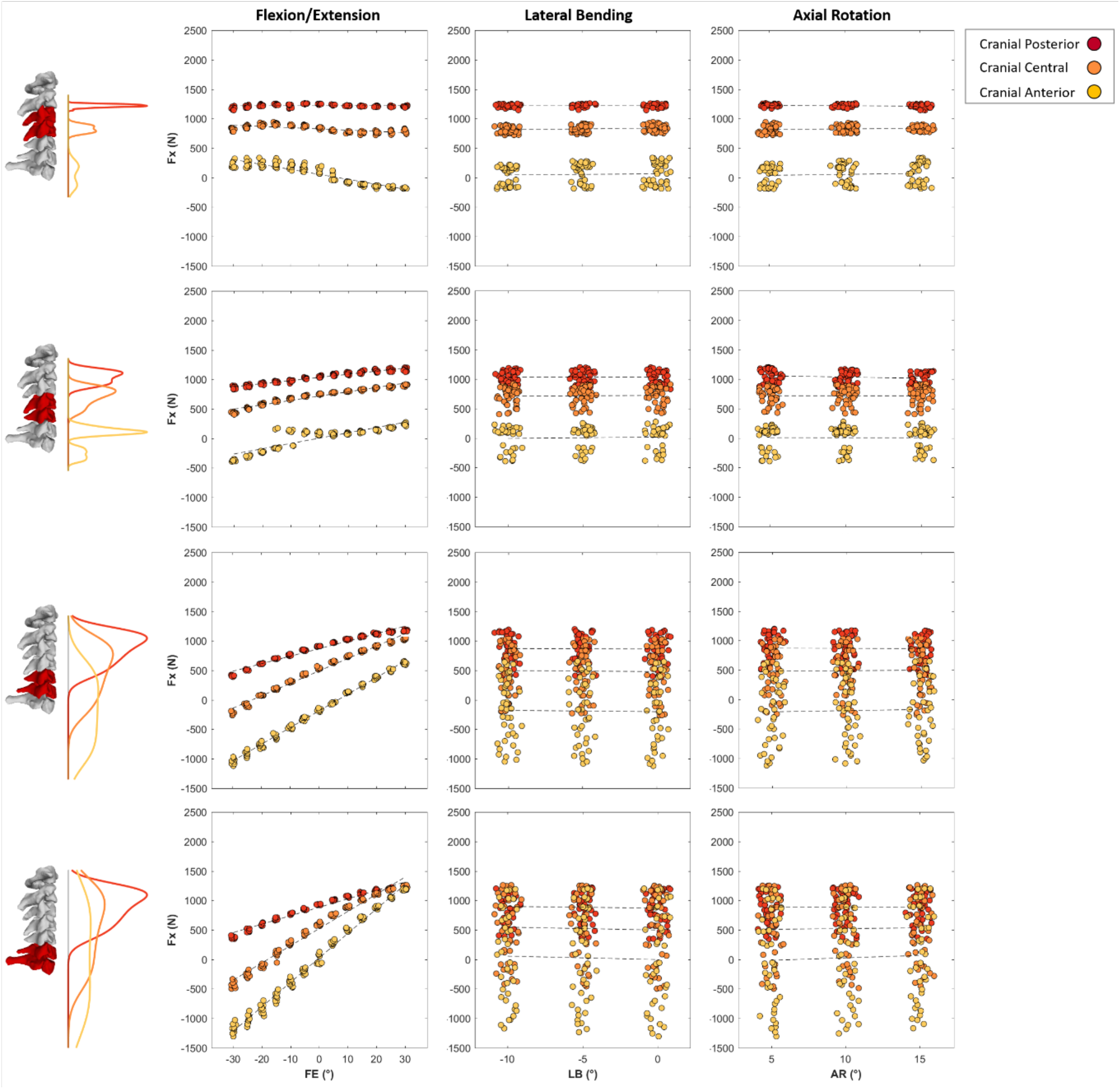
Maximal anteroposterior (anterior (+) and posterior (-) values) shear joint loads (newton) of C3-C4 (top row) to C6-C7 (bottom row) intervertebral joints plotted against 5° changes in Flexion(-)/Extension(+) (left column), Lateral Bending (centre column) and Axial Rotation (right column) during the cranial loading conditions (Cranial Posterior, Cranial Central and Cranial Anterior). Kernel density estimate plots to the left of the subplot rows represent the frequency distribution density of the maximal joint loads on the vertical axes for each loading condition. First order polynomial lines of best fit are plotted to highlight the effect of joint angle on compressive joint loads for each loading condition (dashed lines). In each subplot data points are spread slightly in each 5° bin on the horizontal axes for better visualisation.

**Figure 6.**
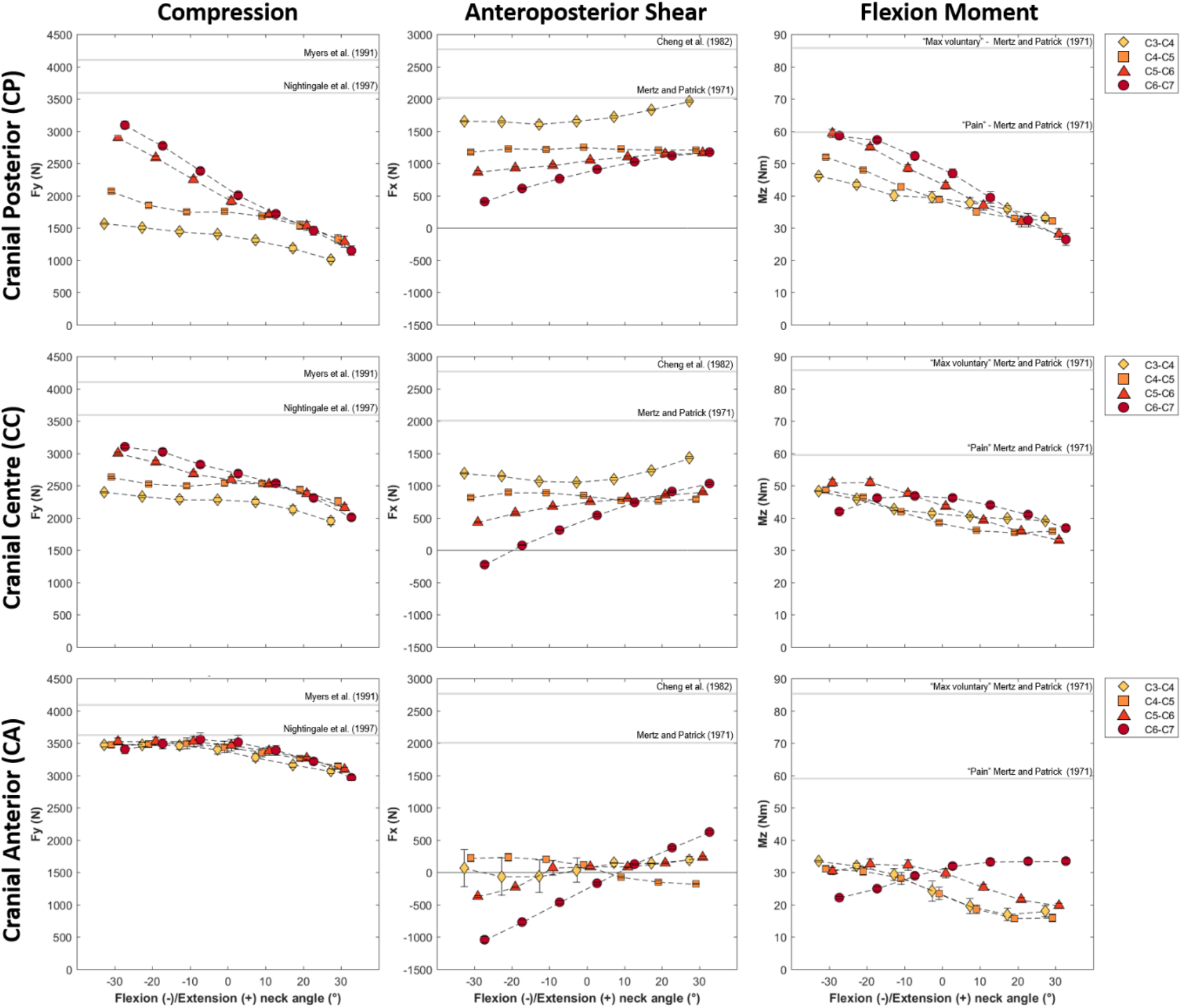
Mean and standard deviation values for maximal compression (left column), anteroposterior (centre column) and flexion moment (right column) of all initial neck angle conditions plotted against changes in neck flexion (negative) and extension (positive) angles for cranial loading conditions (CP – Cranial Posterior, CC – Cranial Central and CA – Cranial Anterior). Estimated injury thresholds from the literature for the entire cervical spine are also presented with the horizontal lines for compression and anteroposterior shear. Flexion moment failure thresholds have been identified to be larger than 150 Nm for the intact cervical spine thus subjective thresholds of “Pain” and “maximum voluntary contraction” are presented

**Figure 7.**
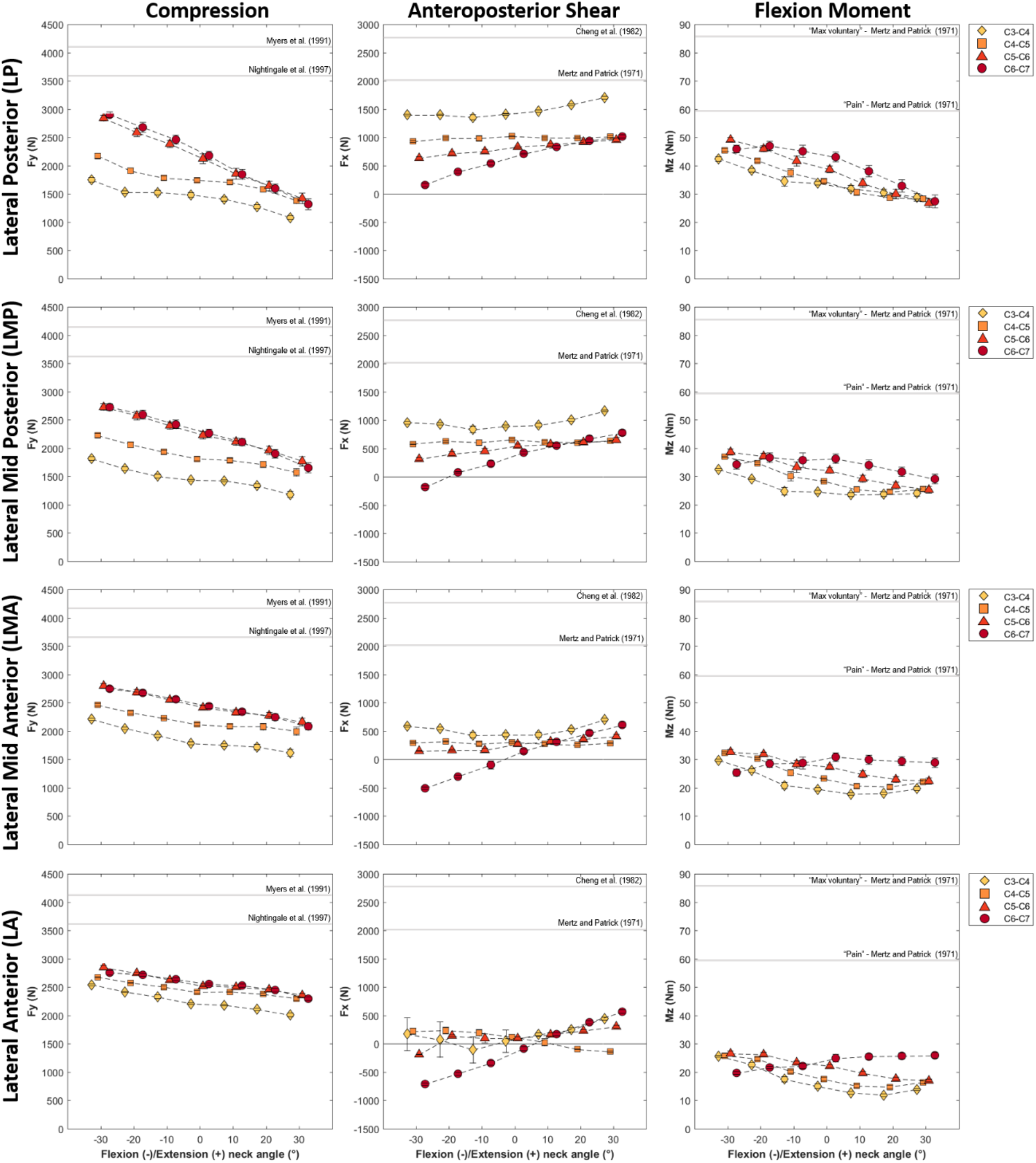
Mean and standard deviation values for maximal compression, anteroposterior (anterior (+) and posterior (-)) and flexion moment of all initial neck angle conditions plotted against changes in neck flexion (negative) and extension (positive) angles for lateral loading conditions (LP – Lateral Posterior, LMP – Lateral Mid-Posterior, LMA – Lateral Mid-Anterior and LA – Lateral Anterior). Estimated injury thresholds from the literature for the entire cervical spine are also presented with the horizontal lines for compression and anteroposterior shear. Flexion moment failure thresholds have been identified to be larger than 150 Nm for the intact cervical spine thus subjective thresholds of “Pain” and “maximum voluntary contraction” are presented.

Lateral bending and axial rotation of the neck did not significantly affect the magnitudes of compressive joint loading across the loading conditions (Figure 4). Maximal compressive joint loads during the 50 ms simulations increased as initial neck position transitioned from an extended (30°) to a flexed position (−30°) with largest loads experienced when the neck was initially flexed. The largest increase was seen in the posterior cranial impacts (CP) during which lower cervical spine compressive loading increased by approximately 50% (from 2100 to 3200 N) in the −30° flexed condition compared to neutral (0°) (Figure 6 – Column 1 Rows 3 and 4). Lateral posterior impacts (LP and LMP) also resulted in increased compressive joint loading of up to 30% (from 2200 to 2900 N). In anterior loading conditions (CA, LMA and LA) initial neck flexion had a smaller effect with compression increasing less than 500 N (~20%) from when the neck was extended (Figure 5 and 6 – Column 1).

Lateral bending and axial rotation of the neck did not affect the magnitudes of anteroposterior loading across the loading conditions (Figure 5). Maximal anteroposterior shear loads changed direction from anterior to posterior as the initial neck flexion angle increased (Figures 6 and 7 – Column 2). This was more evident at the C5-C6 and C6-C7 joint levels and during anterior loading of the skull (CA, LMA and LA), here anterior shear loads of approximately 600 N when the neck was extended changed to posterior loads of 1000 N when it was flexed. Posterior loading conditions (CP, CC, LP and LMP) resulted in anterior shear loading across the initial neck angles and all vertebral joint levels other than C6-C7 in the most flexed conditions (Figures 6 and 7 – Column 2).

Flexion moments increased up to 60 Nm (Figure 6 and 7 – Column 3) as the initial neck flexion angle approached −30°. This flexion moment pattern across neck flexion angles was more visible during the posterior loading conditions (CP and LP). Other loading conditions did not affect neck joint flexion moments across the initial neck angles. Flexion moments were larger in the lower cervical spine when the spine was loaded at the posterior (CP and LP) and across initial neck angles. However, in other loading conditions lower cervical spine flexion moments reduced as neck flexion angles increased.

Lateral shear displayed the lowest joint load magnitudes (< 1000 N). As left lateral bending angle increased left shear loads in the lower cervical spine (C5-C6 and C6-C7) also increased during cranial impacts. Lateral loading conditions (which were on the right side of the skull) increased the left lateral loading. The higher loading rate resulted in larger loads across all initial head angle and loading conditions. Individual values for maximal compressive loading and lateral shear during only cranial impacts are presented in the graphs for brevity. The equivalent graphs for lateral head impacts (LA, LMA, LMP and LP) are available in the supplementary material.

## Discussion

In this study we simulated and examined the dynamic response of the cervical spine to loading conditions representative of accidental head-on rugby tackle injuries using a musculoskeletal modelling approach. The computer simulations were informed by experimental *in vivo* and *in vitro* data representing realistic head-on rugby tackle conditions. The head-on rugby tackles were simulated by applying the impact forces that are normally experienced on the shoulder area to the head, and the effect of tackling technique (neck angle) on cervical spine internal loading was assessed. From an injury mechanisms analysis perspective, the results suggested that hyperflexion is unlikely to be the primary injury mechanisms of the cervical spine in accidental head-on rugby tackling. Three main findings support this conclusion: i) compressive and shear loads were the dominant loading types; ii) shear loads pattern changed across different levels of the cervical spine; iii) peak compressive and shear loads were generated before the neck exceeded its flexion limits. From a tackling technique perspective, the simulation results showed that a higher degree of neck flexion at impact poses the cervical spine in a more hazardous situation and increases the cervical spine compressive and shear loading.

Our computational study expands on the earlier theoretical work by Nightingale, et al. ^28^ by showing that, also during head-on rugby tackles, axial head impacts are likely to be generated by a buckling injury mechanism. This is shown in our simulation results when central and posterior cranial impacts happen in combination with a flexed neck position; in such cases high compression, anterior shear and flexion loads are generated in the lower cervical spine well before the neck approached the anatomical injury limit (45° of flexion)^23^. This supports the hypothesis that anterior facet dislocation, which is the most common catastrophic injury in tackling^13,14^, is generated much earlier than when physiological neck flexion ranges are exceeded loading. In fact, anterior dislocation injuries caused by a hyperflexion of the entire cervical spine have been mostly disregarded ^18,23,52,53^, as *in vitro* experiments have not succeeded in replicating them even under substantial loads (190 Nm of flexion moment). Therefore, our results advise to move away from using “hyperflexion” as the cause of bilateral facet dislocation in the sub-axial cervical spine (C3-C7) under these conditions ^23^. Finally, it is key to underline that buckling might not itself cause injury (tissue failure) but alters pre-injury neck kinematics and resulting loading modalities causing injury in the lower cervical spine (C3-C7)^21^. These alterations in intervertebral loading patterns help explain why injuries cannot be characterised by head motion alone^15,16^, which is an important consideration when relating field injuries to video analysis and players’ recollections of the incident. Computational approach can indeed help to see beyond the external kinematics and should be used to inform injury prevention strategies in the game of Rugby.

Neck flexion angle at the time of impact had the largest effect over lateral bending and axial rotation angles on neck internal loading during head-on tackle simulations. The important role of neck flexion on spinal loading during impacts has been previously shown with *in vitro* and *in silico* experiments^15,28^. Our results confirmed that compressive loading increased with neck flexion also in rugby tackling (Figure 6 and 7), whilst anterior shear was maintained in the mid cervical spine (C3-C4 and C4-C5) or directed posteriorly primarily in the lower cervical spine (C5-C6 and C6-C7) during central and posterior head impacts. A more flexed position causes the neck to lose its natural lordosis resulting in an axial alignment of the vertebrae and a stiffer configuration of the cervical spine. This is a very hazardous situation that alters the transmission of head impact forces through the cervical spine and the way in which the impact energy is dissipated ^28^. Our results supported this theory as compressive joint loads increased with the alignment between the impact force vector and the cervical column axis as neck flexion increased. This is also in line with the experiments reported by Nightingale, et al. ^21^ who showed higher risk for injury when the impact is aligned within 15° of the neck axis. Therefore, we suggest that the main drivers for catastrophic cervical spine injuries during accidental axial head-on rugby impacts are the sagittal plane parameters tested in our study. This is confirmed by the lower effect of out of plane angles (lateral bending and axial rotation) and loading conditions (lateral impacts) on neck loading in our simulations, and supported by previous experimental and theoretical work.

Additionally, the change in relative alignment between the neck and impact force vector is likely to have caused the inverse loading pattern between compression and shear forces observed in the posterior loading conditions in the lower cervical spine (C6-C7). This inverse pattern is observable at an individual joint level, and is characterised by a concurrent increase in compression and decrease in anterior shear forces as neck flexion increases. This inversely proportional effect could be a beneficial compensation between loading modalities when the neck is in a more neutral position. For instance, during posterior cranial impacts (Figure 6, row 1), the proximity to injury thresholds is reduced from 65% to 40% (~ 1000 N) in compression and increased from 17 to 30% in shear loading of shear tolerance values (~ 400 N) when just moving from a 15° extended to a 15° flexed position.

Video analysis has estimated energy transfer during rugby tackle events can vary between 1.4 kJ and 3.0 kJ ^35^ which is considerably more energy than the 82 J needed to cause neck injury *in vitro* ^18^ and in silico ^28^ during axial impacts. This highlights the importance of correct tackling technique to position the tackler’s head away from the oncoming ball carrier to minimise the amount of energy transferred to the neck in an accidental head-on tackle. Low tackles (i.e. targeting the hips and thighs of the ball carrier), which are aimed lateral to the center of mass, are more effective in arresting the ball carrier’s momentum and reduce the possibility of concussion to the tackler, as the tackler’s head is directed away from the ball carrier’s torso-hips. However, this requires the tackler to bend at the waist which could lead to a head, neck and torso alignment if they adopt poor technique (e.g. if fatigued or not wanting to be penalised for a high tackle). Additionally, during a head-on tackle, with the high possibility of head pocketing (i.e. head is constrained by the soft tissue of the abdomen and large impact area with high friction), the tackler’s neck would be required to arrest the momentum of the following body resulting in almost certain catastrophic injury during axial head impacts^15,21,54^. Although challenging to quantify the probability of these pocketing situations future computational and experimental studies could evaluate the effects of the tackler’s head contacting more conforming surfaces of the abdomen, groin or thigh compared to rigid surfaces of bones such as the ribs or pelvis. Head contact to the rigid surfaces that are not covered by as much soft tissue as conforming surfaces may result in more glancing impacts with high head acceleration and risk of concussion^55^. Such a study would provide a more complete overview on the effect of tackle height during accidental head-on tackles with regards to possible concussive head and catastrophic neck injuries.

The important role of active and passive neck muscle forces to load the cervical spine during impacts has been previously investigated ^24–26,28^. Neck musculature provides a compressive preload that increases the stability of the spinal column but also bring the intervertebral loads closer to their critical failure limits ^22^. Our study is the first one in which task specific muscle forces are estimated from simulated *in vivo* rugby tackling and applied to the analysis of sub-axial cervical spine injury mechanisms. Activation levels derived using EMG-assisted methods for the analysis of staged laboratory rugby tackling provided an internal loading condition that would be expected in a tackle where the anticipated impact was to the right shoulder. This approach allowed for the use of physiologically plausible neck muscle forces at the time of impact, as they were not derived from any *a priori* assumptions. Assumed or arbitrary activation levels could result in the overestimation of intervertebral joint loads if maximal joint stiffness was the objective ^29^ and result in initial conditions that are not situation specific. Our proposed approach increased the fidelity of the simulations as neural recruitment strategies during impacts are still not well understood in order to apply explicit *a priori* objective criteria to estimate muscle activations ^46,48^.

Factors associated with the injury severity of a head-on rugby tackle are many^22,55^ and all possible combinations that might be experienced on a rugby field cannot be fully replicated experimentally or computationally. Player internal factors such as experience, physical maturity, fitness and technique ^3,56^ and inciting event characteristics, such as the tackler and ball carrier approach velocities will all influence the risk of vertebral injury. Our study was conducted using an MRI-informed musculoskeletal model of a rugby player which allowed for the prescription of task specific body kinematics and muscle forces. A limitation of our approach is that neck muscle forces and initial angular velocities applied across all conditions were estimated from a single experimental neck position during a tackle. Ideally for each initial neck angle condition simulated in this study an estimation of muscle activations would have been experimentally estimated or optimised based on an *a priori* criterion. However, this was experimentally infeasible and outside the scope of this study. Therefore we would expect that different combination of muscle forces would be present for each neck angle condition. However, the overall magnitude of the pre-compression force produced by the muscles on the spine would not change significantly for an individual positioning their head and neck in different configurations prior to tackling. The chosen loading conditions for our study aimed to replicate impact directions representative of head-on tackles. It should be noted that in reality impacts to the head would result in a shear loading component and translation of the point of force application that would change the resulting impact force vector direction for the duration of the impact event. This is difficult to replicate in multibody models, and validated contact models should be used in the future to examine the effect of tackling technique (e.g. neck and relative torso angles, loading condition) on neck loading with more extensive multibody models. For this reason, it was assumed that a point load would be a reasonable representation for the short durations simulated (50 ms) as has been done in previous musculoskeletal studies^40,57^.

The response of the cervical spine to axial head impacts has been previously shown with simplified finite element models that included rigid vertebrae and bushing elements to represent intervertebral joint behaviour^28^ similar to the present study. Although musculoskeletal models cannot simulate injury, under controlled circumstances they provide a more streamlined solution to run data-driven simulations and yield an initial appreciation of the cervical spine’s response to external impacts. For this reason, the simulations of our study did not continue past the peak external loading of 50 ms as past this point plastic deformation is likely to occur^15^ which cannot be approximated by musculoskeletal models. This is supportive for the use of analytical modelling as patterns of internal loading and the spine’s response can be identified whilst impact parameters, such as neck angle, torso alignment and loading conditions, are programmatically changed. However, to fully understand the cause-effect relationship between the internal loading, resulting kinematic response of the cervical spine, and clinically observed injuries from axial head impacts during accidental head-on rugby tackles, a further step using finite element models should be completed. This step could utilise kinematics (vertebral alignment and joint angular velocities) and muscle forces predicted by the musculoskeletal model, from representative experimental or improved on-field data (e.g. accelerations, quantifiable head/neck/torso angles), as boundary conditions in detailed finite element simulations to identify localised regions of stress and strain on cervical spine structures. These finite element models could provide the specific identifiers of how cervical spine buckling leads to injury on a local vertebral level (e.g. joint dislocation, vertebral fracture, ligament tear or disc burst).

Our study indicates that sub-axial (C3-C7) cervical spine injuries observed in the simulated accidental head-on rugby tackles are likely caused by a loading pattern that is commonly associated to buckling rather than hyperflexion mechanisms. This pattern is characterized by high compression and anterior shear forces, and moderate flexion moment in the mid and lower cervical spine, which is commonly observed in anterior bilateral facet dislocation injuries. Also the simulation results show that a more flexed position of the neck, during impacts on the central and posterior part of the head, increases cervical spine internal loading and therefore the injury risk during head-on rugby impacts. The simulation were guided by experimental data that informed the initial joint angles, angular velocities, muscle forces and external loading conditions providing high external validity to the results. However, the musculoskeletal modelling approach used in our study cannot identify or predict specific types of injury, and finite element models should be integrated to predict vertebral and spinal soft tissues stresses and strain. These findings highlight the importance of the adoption of a correct tackling technique and inclusion of biomechanical analyses to aid in injury prevention strategies and ensure the safety of the athletes in rugby.

## References

1 Brooks, J. H. M., Fuller, C. W., Kemp, S. P. T. & Reddin, D. B. Epidemiology of injuries in English professional rugby union: part 1 match injuries. British Journal of Sports Medicine 39, 757, doi:10.1136/bjsm.2005.018135 (2005).

2 Fuller, C. W. et al. Injury risks associated with tackling in rugby union. British Journal of Sports Medicine 44, 159, doi:10.1136/bjsm.2008.050864 (2010).

3 Tucker, R. et al. Risk factors for head injury events in professional rugby union: a video analysis of 464 head injury events to inform proposed injury prevention strategies. British Journal of Sports Medicine 51, 1152, doi:10.1136/bjsports-2017-097895 (2017).

4 Tierney, G. J., Richter, C., Denvir, K. & Simms, C. K. Could lowering the tackle height in rugby union reduce ball carrier inertial head kinematics? Journal of Biomechanics 72, 29–36, doi:https://doi.org/10.1016/j.jbiomech.2018.02.023 (2018).

5 Tierney, G. J., Denvir, K., Farrell, G. & Simms, C. K. The Effect of Tackler Technique on Head Injury Assessment Risk in Elite Rugby Union. Medicine & Science in Sports & Exercise 50 (2018).

6 Brown, J. C. et al. The incidence of rugby-related catastrophic injuries (including cardiac events) in South Africa from 2008 to 2011: a cohort study. BMJ Open 3, e002475, doi:10.1136/bmjopen-2012-002475 (2013).

7 West, S. W. et al. Trends in match injury risk in professional male rugby union: a 16-season review of 10 851 match injuries in the English Premiership (2002–2019): the Professional Rugby Injury Surveillance Project. British Journal of Sports Medicine 55, 676, doi:10.1136/bjsports-2020-102529 (2021).

8 Fuller, C. W. Catastrophic Injury in Rugby Union. Sports Medicine 38, 975–986, doi:10.2165/00007256-200838120-00002 (2008).

9 Cross, M., Kemp, S., Smith, A., Trewartha, G. & Stokes, K. Professional Rugby Union players have a 60% greater risk of time loss injury after concussion: a 2-season prospective study of clinical outcomes. British Journal of Sports Medicine 50, 926, doi:10.1136/bjsports-2015-094982 (2016).

10 Organization, W. H. & Society, I. S. C. International perspectives on spinal cord injury. Report No. 9241564660, (World Health Organization and International Spinal Cord Society, 2013).

11 Stokes, K. A. et al. Does reducing the height of the tackle through law change in elite men’s rugby union (The Championship, England) reduce the incidence of concussion? A controlled study in 126 games. British Journal of Sports Medicine 55, 220, doi:10.1136/bjsports-2019-101557 (2021).

12 Bahr, R. & Krosshaug, T. Understanding injury mechanisms: a key component of preventing injuries in sport. British Journal of Sports Medicine 39, 24–329, doi:10.1136/bjsm.2005.018341 (2005).

13 Kuster, D., Gibson, A., Abboud, R. & Drew, T. Mechanisms of cervical spine injury in rugby union: a systematic review of the literature. Br J Sports Med 46, 550–554, doi:10.1136/bjsports-2011-090360 (2012).

14 Dennison, C. R., Macri, E. M. & Cripton, P. A. Mechanisms of cervical spine injury in rugby union: is it premature to abandon hyperflexion as the main mechanism underpinning injury? British Journal of Sports Medicine 46, 545 (2012).

15 Nightingale, R. W., McElhaney, J. H., Richardson, W. J. & Myers, B. S. Dynamic responses of the head and cervical spine to axial impact loading. Journal of Biomechanics 29, 307–318, doi:10.1016/0021-9290(95)00056-9 (1996).

16 Yoganandan, N., Pintar, F. A., Sances, A., Jr., Reinartz, J. & Larson, S. J. Strength and Kinematic Response of Dynamic Cervical Spine Injuries. Spine 16 (1991).

17 Badenhorst, M., Verhagen, E., Lambert, M. I., van Mechelen, W. & Brown, J. C. ‘In a blink of an eye your life can change’: experiences of players sustaining a rugby-related acute spinal cord injury. Injury Prevention 25, 313–320, doi:10.1136/injuryprev-2018-042871 (2019).

18 Nightingale, R. W., Doherty, B. J., Myers, B. S., McElhaney, J. H. & Richardson, W. J. (SAE International, 1991).

19 Ivancic, P. C. Cervical spine instability following axial compression injury: A biomechanical study. Orthopaedics & Traumatology: Surgery & Research 100, 127–133, doi:https://doi.org/10.1016/j.otsr.2013.10.015 (2014).

20 Ivancic, P. C. Biomechanics of Sports-Induced Axial-Compression Injuries of the Neck. Journal of Athletic Training 47, 489–497, doi:10.4085/1062-6050-47.4.06 (2012).

21 Nightingale, R. W. et al. The Dynamic Responses of the Cervical Spine: Buckling, End Conditions, and Tolerance in Compressive Impacts. SAE International 106, 3968–3988, doi:10.4271/973344 (1997).

22 Saari, A. et al. Compressive Follower Load Influences Cervical Spine Kinematics and Kinetics During Simulated Head-First Impact in an in Vitro Model. Journal of Biomechanical Engineering 135, 111003-111003-111011, doi:10.1115/1.4024822 (2013).

23 Nightingale, R. W., Bass, C. R. & Myers, B. S. On the relative importance of bending and compression in cervical spine bilateral facet dislocation. Clinical Biomechanics 64, 90–97, doi:https://doi.org/10.1016/j.clinbiomech.2018.02.015 (2019).

24 Dibb, A. T. et al. Importance of Muscle Activations for Biofidelic Pediatric Neck Response in Computational Models. Traffic Injury Prevention 14, 116–127, doi:10.1080/15389588.2013.806795 (2013).

25 Happee, R., de Bruijn, E., Forbes, P. A. & van der Helm, F. C. T. Dynamic head-neck stabilization and modulation with perturbation bandwidth investigated using a multisegment neuromuscular model. Journal of Biomechanics 58, 203–211, doi:10.1016/j.jbiomech.2017.05.005 (2017).

26 de Bruijn, E., van der Helm, F. C. T. & Happee, R. Analysis of isometric cervical strength with a nonlinear musculoskeletal model with 48 degrees of freedom. Multibody System Dynamics 36, 339–362, doi:10.1007/s11044-015-9461-z (2016).

27 Chancey, V. C., Nightingale, R. W., Van Ee, C. A., Knaub, K. E. & Myers, B. S. Improved estimation of human neck tensile tolerance: reducing the range of reported tolerance using anthropometrically correct muscles and optimized physiologic initial conditions. Stapp Car Crash J 47, 135–153 (2003).

28 Nightingale, R. W., Sganga, J., Cutcliffe, H. & Bass, C. R. Impact responses of the cervical spine: A computational study of the effects of muscle activity, torso constraint, and pre-flexion. J Biomech 49, 558–564, doi:10.1016/j.jbiomech.2016.01.006 (2016).

29 Mortensen, J., Trkov, M. & Merryweather, A. Exploring novel objective functions for simulating muscle coactivation in the neck. Journal of Biomechanics 71, 127–134, doi:https://doi.org/10.1016/j.jbiomech.2018.01.030 (2018).

30 Seminati, E., Cazzola, D., Preatoni, E. & Trewartha, G. Specific tackling situations affect the biomechanical demands experienced by rugby union players. Sports Biomechanics 16, 58–75, doi:10.1080/14763141.2016.1194453 (2017).

31 Cazzola, D., Holsgrove, T. P., Preatoni, E., Gill, H. S. & Trewartha, G. Cervical Spine Injuries: A Whole-Body Musculoskeletal Model for the Analysis of Spinal Loading. PLoS One 12, e0169329, doi:10.1371/journal.pone.0169329 (2017).

32 Lloyd, D. G. & Besier, T. F. An EMG-driven musculoskeletal model to estimate muscle forces and knee joint moments in vivo. Journal of Biomechanics 36, 765–776, doi:https://doi.org/10.1016/S0021-9290(03)00010-1 (2003).

33 Silvestros, P. et al. Electromyography-Assisted Neuromusculoskeletal Models Can Estimate Physiological Muscle Activations and Joint Moments Across the Neck Before Impacts. Journal of Biomechanical Engineering, doi:10.1115/1.4052555 (2021).

34 Seminati, E., Preatoni, E., Trewartha, G., Wallbaum, A. & Cazzola, D. in Abstract Book of the 26th Congress of the International Society of Biomechanics, Brisbane (Aus), July 23–27, 2017.

35 Hendricks, S., Karpul, D. & Lambert, M. Momentum and kinetic energy before the tackle in rugby union. Journal of sports science & medicine 13, 557 (2014).

36 Delp, S. L. et al. OpenSim: Open-Source Software to Create and Analyze Dynamic Simulations of Movement. IEEE Transactions on Biomedical Engineering 54, 1940–1950, doi:10.1109/TBME.2007.901024 (2007).

37 Mortensen, J. D., Vasavada, A. N. & Merryweather, A. S. The inclusion of hyoid muscles improve moment generating capacity and dynamic simulations in musculoskeletal models of the head and neck. PLOS ONE 13, e0199912, doi:10.1371/journal.pone.0199912 (2018).

38 Vasavada, A. N., Li, S. P. & Delp, S. L. Influence of muscle morphometry and moment arms on the moment-generating capacity of human neck muscles. Spine 23, 412–422, doi:10.1097/00007632-199802150-00002 (1998).

39 Silvestros, P. et al. Musculoskeletal modelling of the human cervical spine for the investigation of injury mechanisms during axial impacts. PLOS ONE 14, e0216663, doi:10.1371/journal.pone.0216663 (2019).

40 Kuo, C. et al. Passive cervical spine ligaments provide stability during head impacts. Journal of The Royal Society Interface 16, 20190086, doi:10.1098/rsif.2019.0086 (2019).

41 Kawasaki, T. et al. Kinematics of Rugby Tackling: A Pilot Study With 3-dimensional Motion Analysis. The American Journal of Sports Medicine 46, 2514–2520, doi:10.1177/0363546518781808 (2018).

42 Gianotti, S. M., Quarrie, K. L. & Hume, P. A. Evaluation of RugbySmart: A rugby union community injury prevention programme. Journal of Science and Medicine in Sport 12, 371–375, doi:https://doi.org/10.1016/j.jsams.2008.01.002 (2009).

43 Quarrie, K. et al. RugbySmart: Challenges and Lessons from the Implementation of a Nationwide Sports Injury Prevention Partnership Programme. Sports Medicine 50, 227–230, doi:10.1007/s40279-019-01177-8 (2020).

44 Pizzolato, C. et al. CEINMS: A toolbox to investigate the influence of different neural control solutions on the prediction of muscle excitation and joint moments during dynamic motor tasks. Journal of Biomechanics 48, 3929–3936, doi:https://doi.org/10.1016/j.jbiomech.2015.09.021 (2015).

45 Sartori, M., Farina, D. & Lloyd, D. G. Hybrid neuromusculoskeletal modeling to best track joint moments using a balance between muscle excitations derived from electromyograms and optimization. Journal of Biomechanics 47, 3613–3621, doi:https://doi.org/10.1016/j.jbiomech.2014.10.009 (2014).

46 Fice, J. B., Siegmund, G. P. & Blouin, J.-S. Neck muscle biomechanics and neural control. Journal of Neurophysiology 120, 361–371, doi:10.1152/jn.00512.2017 (2018).

47 Newell, R. S., Blouin, J.-S., Street, J., Cripton, P. A. & Siegmund, G. P. The neutral posture of the cervical spine is not unique in human subjects. Journal of Biomechanics 80, 53–62, doi:https://doi.org/10.1016/j.jbiomech.2018.08.012 (2018).

48 Siegmund, G. P., Blouin, J.-S., Carpenter, M. G., Brault, J. R. & Inglis, J. T. Are cervical multifidus muscles active during whiplash and startle? An initial experimental study. BMC Musculoskeletal Disorders 9, 80–80, doi:10.1186/1471-2474-9-80 (2008).

49 Race, D. A., Broom, D. N. & Robertson, D. P. Effect of Loading Rate and Hydration on the Mechanical Properties of the Disc. Spine 25, 662–669 (2000).

50 Newell, N., Grigoriadis, G., Christou, A., Carpanen, D. & Masouros, S. D. Material properties of bovine intervertebral discs across strain rates. Journal of the Mechanical Behavior of Biomedical Materials 65, 824–830, doi:https://doi.org/10.1016/j.jmbbm.2016.10.012 (2017).

51 Tanabe, Y. et al. The kinematics of 1-on-1 rugby tackling: a study using 3-dimensional motion analysis. Journal of Shoulder and Elbow Surgery 28, 149–157, doi:https://doi.org/10.1016/j.jse.2018.06.023 (2019).

52 Mertz, H. J. & Patrick, L. M. Strength and Response of the Human Neck. SAE International, doi:https://doi.org/10.4271/710855 (1971).

53 Mertz, H. J. & Patrick, L. M. Investigation of the Kinematics and Kinetics of Whiplash. SAE International, doi:https://doi.org/10.4271/670919 (1967).

54 Camacho, D. L. A., Nightingale, R. W. & Myers, B. S. The Influence of Surface Padding Properties on Head and Neck Injury Risk. Journal of Biomechanical Engineering 123, 432–439, doi:10.1115/1.1389086 (2001).

55 Nightingale, R. W., McElhaney, J. H., Richardson, W. J., Best, T. M. & Myers, B. S. Experimental Impact Injury to the Cervical Spine: Relating Motion of the Head and the Mechanism of Injury. JBJS 78 (1996).

56 Quarrie, K. L., Cantu, R. C. & Chalmers, D. J. Rugby Union Injuries to the Cervical Spine and Spinal Cord. Sports Medicine 32, 633–653, doi:10.2165/00007256-200232100-00003 (2002).

57 Mortensen, J. D., Vasavada, A. N. & Merryweather, A. S. Sensitivity analysis of muscle properties and impact parameters on head injury risk in american football. Journal of Biomechanics, 109411, doi:https://doi.org/10.1016/j.jbiomech.2019.109411 (2020).

