## Supplementary Figures for "Hyperflexion is unlikely to be the primary cervical spine injury mechanism in accidental head-on rugby tackling"

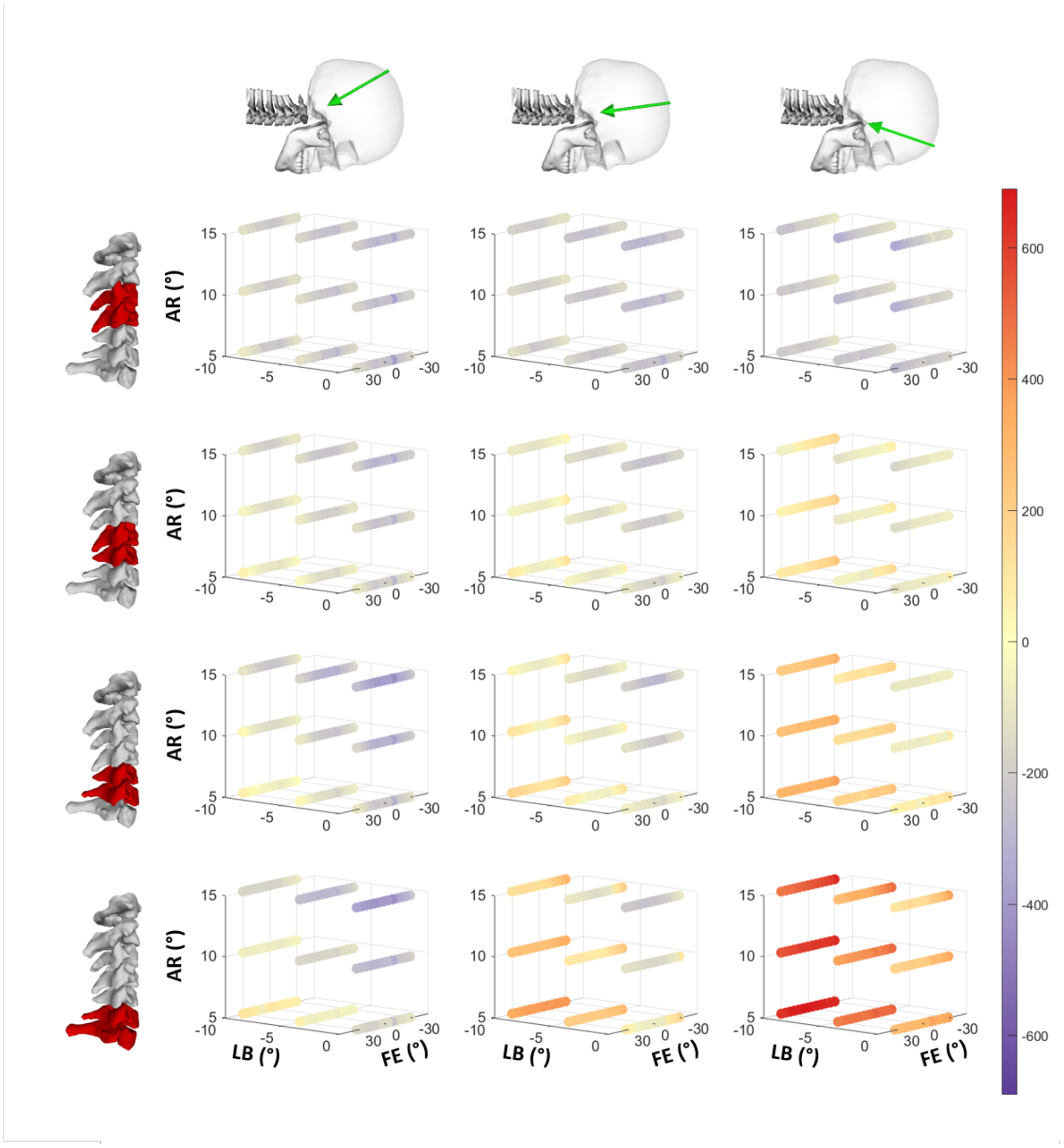


**Figure 1.** Overview presentation of the effects of neck angle (FE = flexion/extension, LB = lateral bending, and AR = axial rotation), external cranial loading location (posterior (CP), central (CC) and anterior (CA)) on maximal cervical spine loading across the cervical spine. Patterns of lateral shear (Fz) loading in Newton sustained during the 50 ms simulations across all simulated initial neck angles for cranial loading conditions. Column represents an individual loading condition (Cranial Posterior – left columns; Cranial Central – centre columns and Cranial Anterior – right columns). Rows represent the cervical spine levels from C3-C4 (top) to C6-C7 (bottom). The cubic grids of each subplot represent the initial neck angle (°) in Flexion/Extension (FE), Lateral Bending (LB) and Axial Rotation (AR). Magnitude of maximal loading (newton) in the 50 ms simulations is represented with the colour bars. Note compression are only positive values and anteroposterior shear positive and negative values to represent direction with anterior and posterior respectively.

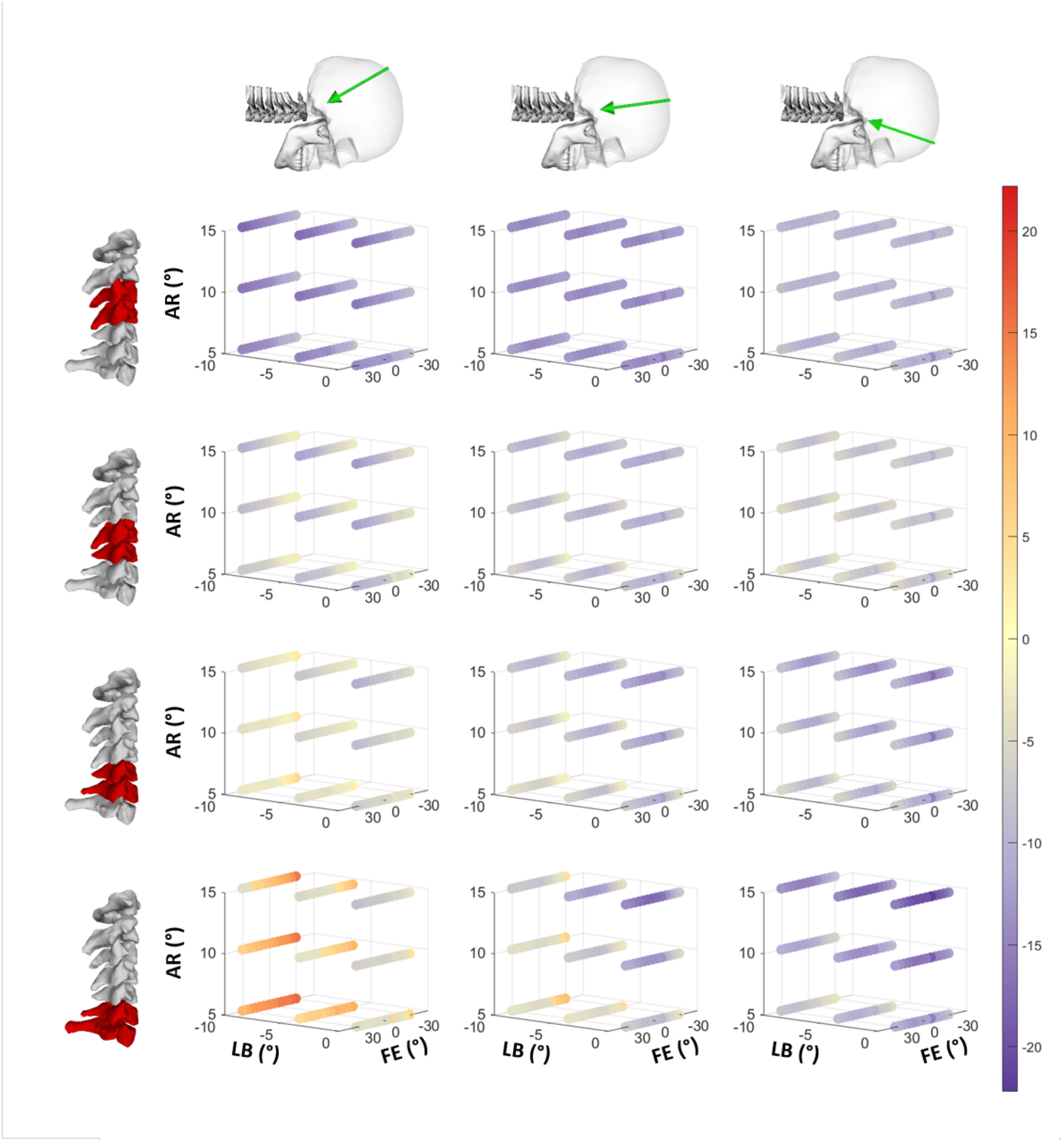


**Figure 2.** Overview presentation of the effects of neck angle (FE = flexion/extension, LB = lateral bending, and AR = axial rotation), external cranial loading location (posterior (CP), central (CC) and anterior (CA)) on maximal cervical spine loading across the cervical spine. Patterns of lateral bending moment (Mx) loading in Nm sustained during the 50 ms simulations across all simulated initial neck angles for cranial loading conditions. Column represents an individual loading condition (Cranial Posterior – left columns; Cranial Central – centre columns and Cranial Anterior – right columns). Rows represent the cervical spine levels from C3-C4 (top) to C6-C7 (bottom). The cubic grids of each subplot represent the initial neck angle (°) in Flexion/Extension (FE), Lateral Bending (LB) and Axial Rotation (AR). Magnitude of maximal loading (newton) in the 50 ms simulations is represented with the colour bars. Note compression are only positive values and anteroposterior shear positive and negative values to represent direction with anterior and posterior respectively.

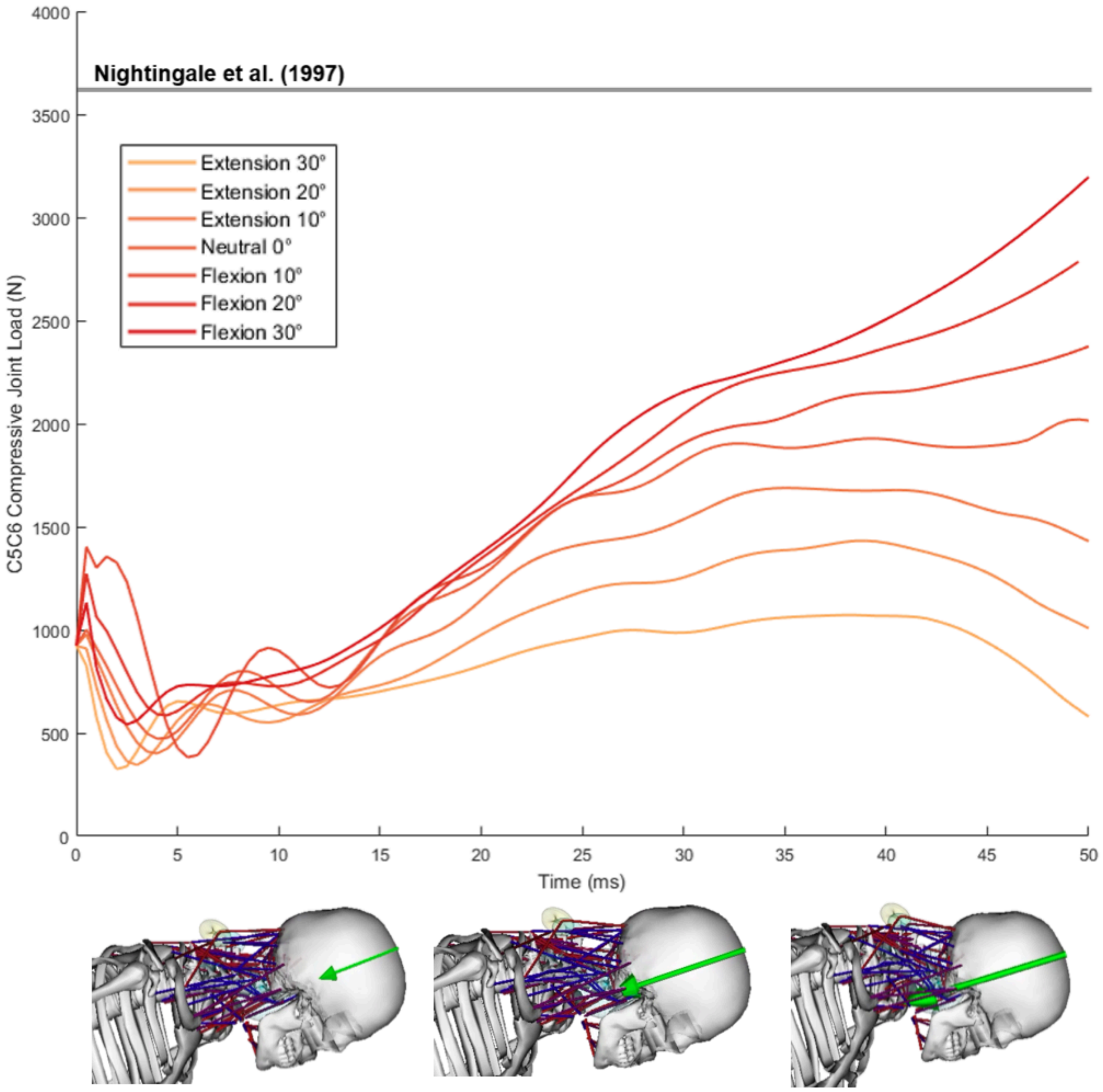


**Figure 3.** Compressive C5-C6 joint load (N) curves are shown for 7 different neck flexion angles (-30° extension to 30° flexion, 0° lateral bending, and 5° axial rotation). Each curve represents the C5-C6 joint reaction force during a single simulation in which the neck muscles generated force using the same activations, and the same force was applied in the cranial central position. Impact occurs at 0 milliseconds (ms). A compressive injury threshold17 is shown for reference (horizontal solid grey line). A visual representation of the simulation is included at the bottom of the graph, where three simulation frames show the different position of the cervical spine and load applied (green arrow) across the first 50 ms.

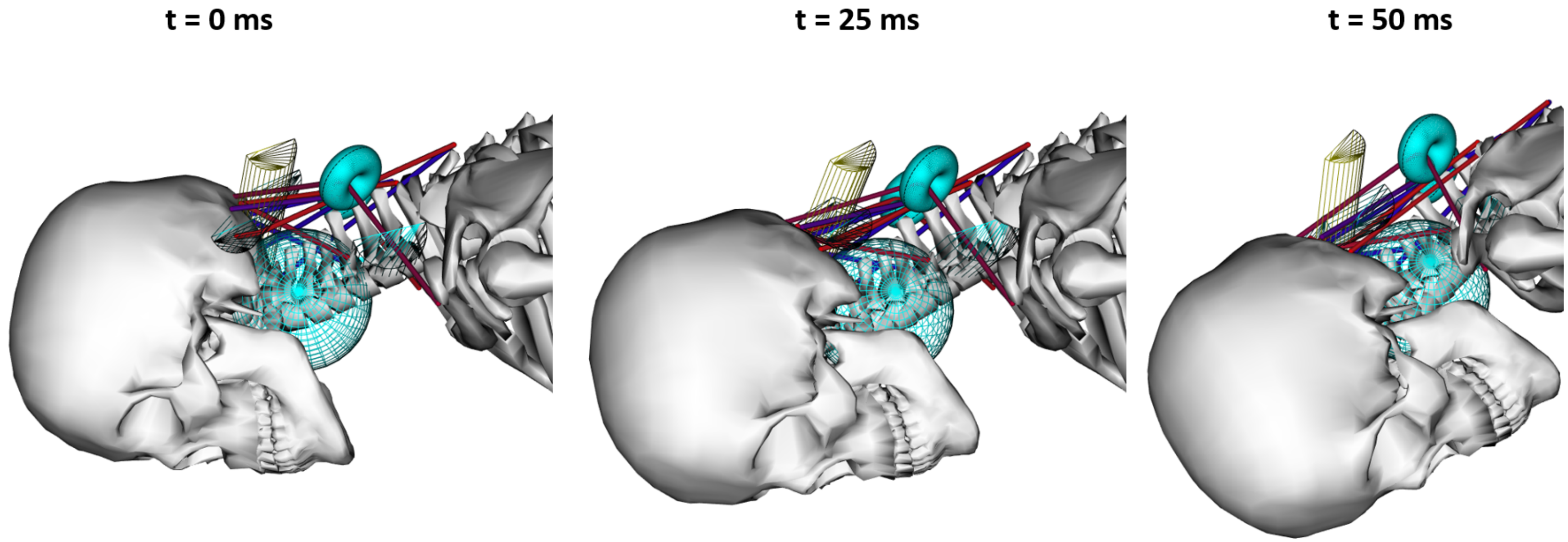


**Figure 4.** Snap shots at the start (0 ms), mid (25 ms) and end (50 ms) of a 50 ms simulation of a Cranial Posterior impact with an initial neck flexion angle of 10 degrees. No hyperflexion of the cervical spine is observed.
